# Short-term and bystander effects of radiation on murine submandibular glands

**DOI:** 10.1101/2022.03.20.484987

**Authors:** Hitoshi Uchida, Matthew H. Ingalls, Eri O. Maruyama, Carl J. Johnston, Eric Hernady, Roberta C. Faustoferri, Catherine E. Ovitt

## Abstract

Many patients treated for head and neck cancers experience salivary gland hypofunction due to radiation damage. Understanding the mechanisms of cellular damage induced by radiation treatment is important in order to design methods of radioprotection. In addition, it is crucial to recognize the indirect effects of IR and the systemic responses that may alter saliva secretion. In this study, radiation was delivered to murine submandibular glands (SMG) bilaterally, using a ^137^Cs gamma ray irradiator, or unilaterally, using a small animal radiation research platform (SARRP). Analysis at 3, 24 and 48 hours showed dynamic changes in mRNA levels and protein abundance in SMG irradiated bilaterally. Unilateral irradiation using the SARRP caused similar changes in the irradiated SMG, as well as significant off-target, bystander effects in the non-irradiated contralateral SMG.

**Summary statement:** Rapid changes in gene expression occur in irradiated salivary glands within hours of irradiation, and in non-irradiated contralateral glands, due to bystander effects.

## Introduction

Over 800,000 new cases of head and neck cancers are diagnosed annually in the world and are treated with a combination of radiation, chemotherapy, and surgery (Cramer et al., 2019). A consequence of radiotherapy for head and neck cancer is hyposalivation, manifested as a significant reduction in saliva flow and permanent loss of the secretory acinar cells (Pinna et al., 2015). Hyposalivation can cause burning mouth, dental caries, gingivitis, periodontitis, and oral infections, as well as difficulty in speaking, chewing, and swallowing, reducing quality of life. Available treatments are only temporary and palliative (Vissink et al., 2010; Villa et al., 2015).

The slow turnover of salivary gland acinar cells is inconsistent with their acute sensitivity to irradiation (IR) (Vissink et al., 2015). The response of salivary glands to IR has been divided into two stages (Coppes et al., 2001; Jasmer et al., 2020). The first stage includes short-term effects that occur within hours to days, such as acute reduction in salivation, changes in saliva composition, interstitial edema and enlarged acinar cells (Coppes et al., 2001). The second stage involves long-term and irreversible effects manifested after weeks to months, including acinar cell loss, fibrosis, continued hyposalivation and absence of cell renewal (Coppes et al., 2001; Konings et al., 2005). The molecular mechanisms driving these changes are not well understood. In addition to direct effects, radiation also induces responses in non-irradiated tissues, known as bystander effects (Blyth and Sykes 2011; Daguenet et al. 2020). In minipigs, the irradiated and non-irradiated, contralateral salivary glands showed a coupled response (Lombaert et al., 2020), but bystander effects in salivary glands have not yet been carefully investigated.

In this study, we utilized a Shepherd Marl I ^137^Cs gamma ray irradiator, with anatomical targeting (achieved using a slit collimator set-up) to deliver radiation bilaterally to murine submandibular glands (SMG). Alternatively, we targeted radiation unilaterally to a single SMG using CT-image guidance with a small animal X-irradiator (Small Animal Radiation Research Platform, SARRP; XStrahl Inc., Suwanee, GA). Use of the SARRP ensured the necessary level of precision for the unilaterally-targeted SMG irradiations. Following unilateral or bilateral irradiation, analysis of mRNA expression and protein abundance showed similar, time-dependent changes in SMGs. In addition, comparison of the irradiated to the contralateral (CL) SMGs revealed significant off-target, bystander effects in the non-irradiated SMG.

## Results

### Short- and long-term loss of saliva secretion following irradiation

To trace the rate of secretory loss in murine salivary glands following IR, we measured saliva flow at 3, 24 and 48 hours, and weekly for up to 12 weeks (Fig. S1G). The baseline saliva flow was established for each mouse on 3 consecutive days, and was determined to be 8.59 ± 1.80 mg/g body weight (Fig. S1H, I; n = 11 - 13). Half the mice received a radiation dose of 15 Gy simultaneously to both left and right SMGs using a ^137^Cs gamma ray irradiator with a slit collimator. At 3 and 48 hours, secretion levels from IR-treated mice were significantly lower than baseline measurements (Fig. S1I). Notably, saliva flow was recovered at 24 hours in both IR-treated and non-IR-treated groups. Although the drop was greater in IR-treated mice, saliva flow decreased again in both IR-treated and non-IR-treated groups by 48 hours. A similar recovery was observed in all mice between the 48-hour and 2-week collection points. The pattern of transient recovery suggests that secretion loss is exacerbated by repeated administration of ketamine.

Saliva volume from non-IR-treated mice did not show significant changes over the 13-week timeline (Fig. S1H), whereas a progressive decrease in saliva secreted from IR-treated mice was measured between 3 and 9 weeks, which leveled off by 12 weeks (Fig. S1I). These data offer important information for the development of intervention or regenerative strategies to prevent salivary gland hypofunction.

### Radiation perturbs expression of tight junction proteins

At early time points following irradiation, changes in cell size and interstitial space consistent with edema were observed in sections of SMG (Fig. S2). We examined whether these changes correlated with altered expression of molecules in the epithelial barrier, including the tight junction proteins ZO-1, Claudin-3 and Claudin-4. In murine SMG, Claudin-3 is expressed by acinar cells, while Claudin-4 is predominantly localized to duct cells (Baker, 2016; Zhang et al., 2018). There was no significant change in ZO-1 immunostaining (Fig. 1A-H) or *Zo-1* mRNA expression levels (Fig. 1I) at 3-, 24- or 48 hours post-IR in mice irradiated with the Cs source. Claudin-3 (CLN3) protein staining was not altered (Fig. 1A-D); however, an increase in Claudin-4 (CLN4) staining was observed (Fig. 1F-H arrowheads) at the basal surface of duct cells, compared to the control salivary glands (Fig. 1E, arrows), which express CLN4 only at the apical surface. A significant increase in both *Claudin-3* and *Claudin-4* mRNA was detected at 3 hours post-IR (Fig. 1J, K). Consistent with previous findings (Yokoyama et al., 2017), CLN3 and CLN4 proteins were increased in abundance on western blots (Fig. 1L-N).

**Fig. 1.**
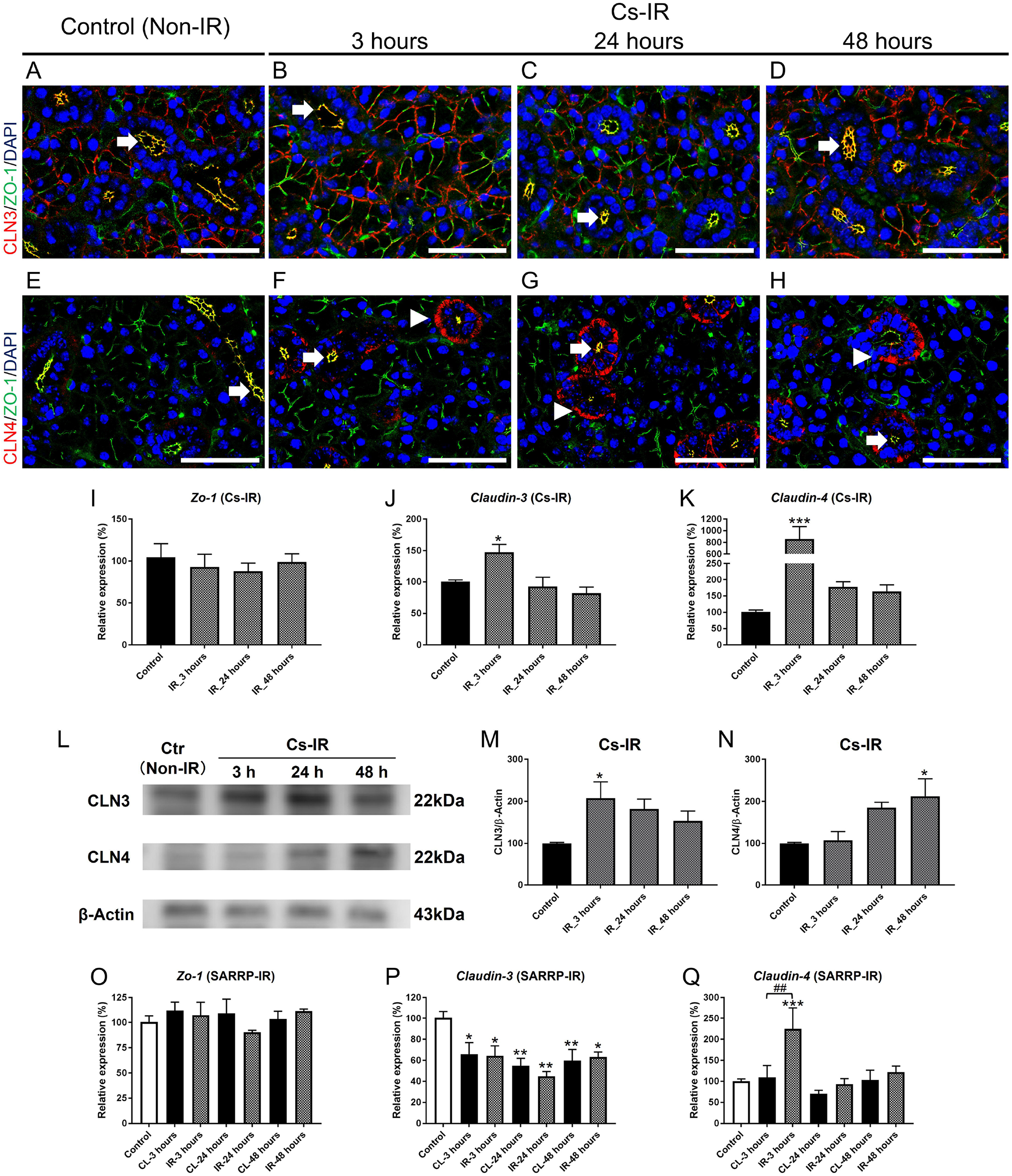
Radiation alters expression of tight junction proteins. **A-D**, Immunofluorescent staining for ZO-1 (green) and CLN3 (red) shows expression on acinar cell membranes, and co-localization at the luminal surface of duct cells (arrows) in control (**A**) and irradiated SMGs at 3 hours (**B**), 24 hours (**C**) and 48 hours (**D**) post-IR. **E-H**, IHC for ZO-1 (green) and CLN4 (red) shows co-localization at the duct cell luminal surface (arrows) in control (**E**) and increased expression of CLN4 (arrowheads) at basal surface of duct cells in irradiated SMGs at 3 hours (**F**), 24 hours (**G**) and 48 hours (**H**) post-IR. Scale bars = 50μm. **I**, Expression level of *Zo-1* mRNA did not change after IR from the Cs source (p > 0.05 vs. control for all time points). **J**,**K** *Claudin-3* (**J**) and *Claudin-4* (**K**) mRNAs were up-regulated at 3 hours post-IR (*Claudin-3*: * p = 0.018 [3 hours]; *Claudin-4*: * p < 0.001 [3 hours]). **L**, Western blots to detect CLN3, CLN4 and β-Actin proteins in control and irradiated SMGs using Cs source (n = 3). **M**,**N**, Quantification of CLN3 and CLN4 proteins detected on western blots. (CLN-3: * p = 0.037 [3 hours]; CLN4: * p = 0.027 [48 hours]). **O**, *Zo-1* mRNA expression did not change in IR or contralateral (CL) SMGs following unilateral IR using SARRP (one-way ANOVA; F = 0.758; two-way ANOVA; F = 0.728, p > 0.05). **P**, In contrast, *Claudin-3* mRNA was down regulated in IR and CL SMGs up to 48 hours post-IR. **Q**, *Claudin-4* mRNA was significantly increased in IR, but not in CL SMGs at 3 hours post-IR (##p < 0.01). Error bars indicate mean ± SEM. Statistical analysis was performed compared to control (non-IR) using one-way ANOVA with Dunnett’s post-hoc test, or comparing IR and CL SMG, using two-way ANOVA with Bonferroni test (n = 3-5 each group).

After unilateral irradiation using the SARRP, *Zo-1* and *Claudin-4* mRNA expression patterns were similar to those in bilaterally irradiated SMG (Fig. 1O, Q). *Claudin-4* mRNA was elevated in the IR-treated, but not in the non-treated contralateral (CL) SMG (Fig.1Q). However, in contrast to the upregulation seen with the Cs source (Fig.1J), *Claudin-3* mRNA expression was reduced in both SARRP-irradiated and CL SMGs (Fig.1P). These results indicate that cell-cell junctions are perturbed both directly and indirectly after irradiation.

### Radiation transiently down-regulates expression of genes involved in saliva secretion

To determine whether radiation disrupts proteins involved in saliva secretion, we investigated the expression profiles and localization of the water channel aquaporin 5 (Aqp5) and the muscarinic receptor type 3 (M3r), which transmits signals from the parasympathetic nervous system. M3R and MIST1, an acinar cell-specific transcription factor (Pin et al., 2000), are co-localized in acinar cells. Immunostaining showed no detectable change in MIST1, M3R or AQP5 (Choi et al., 2009) protein staining at 3, 24, or 48 hours after IR (Fig. 2A-H).

**Fig. 2.**
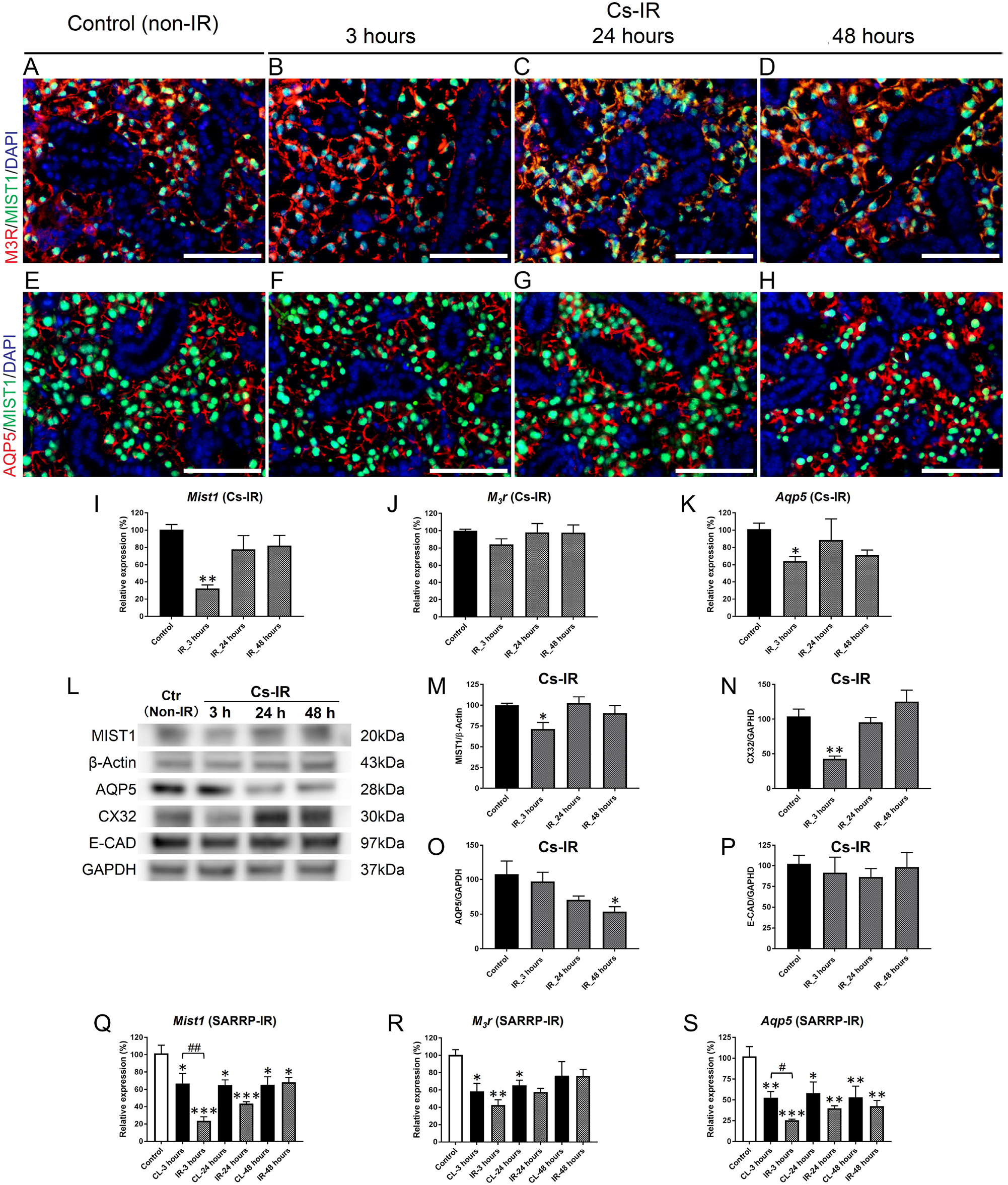
Radiation transiently down-regulates expression of genes involved in saliva secretion. **A-D**, Immunofluorescent staining shows co-localization of M3R (red) with MIST1-positive (green) acinar cells **(A**) on control. M3R expression does not change at (**B**) 3 hours, (**C**) 24 hours, (**D**) 48 hours after IR using the Cs source. **E-H**, AQP5 (red) localization at the apical surface of MIST1-positive (green) acinar cells in control (**E**) does not changes at (**F**) 3 hours, (**G**) 24 hours, (**H**) 48 hours post-IR. Scale bars = 100μm. **I**, Expression of *Mist1* mRNA was significantly reduced at 3 hours post-IR (** indicates p = 0.002), but recovered by 24 hours post-IR (p >0.1). **J**, Relative expression of *M*_*3*_*r* mRNA did not change post-IR using Cs source irradiator (p > 0.1). **K**, *Aqp5* mRNA expression was decreased at 3 hours post-IR (* indicates p = 0.023), but recovered by 24 hours post-IR (p > 0.05). **L**, Western blot of MIST1, AQP5, CX32, E-cadherin (E-CAD), GAPDH and β-Actin proteins in control and SMGs irradiated using Cs source (n = 3). **M**, MIST1 and **N**, CX32 were decreased at 3 hours post-IR (MIST1: * p = 0.041; CX32: **p = 0.003). **O**, AQP5 protein levels declined through 48 hours post-IR (* p <0.05). **P**, In contrast, the level of E-CAD protein was not altered post-IR (one-way ANOVA; F = 0.215, p = 0.883). **Q**, Following SARRP irradiation, relative expression of *Mist1* mRNA was significantly decreased in both IR and CL SMG at 3 hours (***p < 0.001, *p < 0.05, respectively), at 24 hours (***p < 0.001, *p < 0.05, respectively) and at 48 hours (*p < 0.05) vs. control. *Mist1* expression was significantly lower in the IR-treated SMG compared to the CL SMG (##p < 0.01). **R**, Irradiation using SARRP also decreased the relative expression of *M*_*3*_*r* mRNA in both IR and CL SMG at 3 hours (**p < 0.001, *p < 0.01, respectively) vs. non-IR control. **S**, Following SARRP irradiation, relative expression of *Aqp5* mRNA was decreased in both IR and CL SMG at 3 hours vs. control (***p < 0.001, p < 0.01, respectively); at 24 hours (**p < 0.01, *p < 0.05) and at 48 hours (**p < 0.01, **p < 0.01), respectively. *Aqp5* expression was significantly lower in the IR-treated SMG compared to the CL SMG (#p < 0.05). Error bars indicate mean ± SEM. Statistical analysis was performed compared to control (non-IR) using one-way ANOVA with Dunnett’s post-hoc test, or comparing IR and CL SMG using two-way ANOVA with Bonferroni test (n = 3-5 each group).

However, *Mist1* mRNA expression was significantly decreased by 3 hours post-IR, and recovered by 24 hours (Fig. 2I). There was no change in *M3r* mRNA levels (Fig. 2J), but *Aqp5* mRNA expression was down-regulated in IR SMGs at 3 hours post-IR (Fig. 2K). Like *Mist1, Aqp5* mRNA expression was recovered by 24 hours (Fig. 2K). Western blots showed decreases in MIST1 (Fig. 2L, M) and Connexin 32 (the gap junction protein, CX32) (Fig. 2L, N) protein abundance at 3 hours post-IR, but both were recovered by 24 hours. AQP5 protein levels were decreased by 48 hours post-IR (Fig. 2L, O), consistent with an earlier report (Choi et al., 2009). As a control, we quantified E-CAD, a cell adhesion protein, and found that the relative amount did not change within 48 hours post-IR (Fig. 2L, P), as previously reported (Wong et al., 2018). The transient decrease in MIST1, CX32 and AQP5 expression at 3 hours post-IR correlates with the initial drop in saliva secretion.

The expression profiles of *Mist1, M3r* and *Aqp5* mRNA responded similarly in SARRP-irradiated SMGs (Fig. 2Q-S). Notably, expression of these mRNAs also decreased in the CL SMGs, in comparison to non-IR controls, indicating that gene expression is perturbed through both direct and indirect bystander effects.

### DNA damage and inflammatory factors are rapidly up-regulated following irradiation

To assess the level of DNA damage at the 3-, 24- and 48 hour time points following IR with the Cs source, we co-stained SMG sections for γH2AX, which labels radiation-induced double-stranded DNA breaks (Löbrich et al., 2010), and for the sodium/potassium/two-chloride channel (NKCC1), an acinar cell marker. The number of γH2AX foci was significantly increased in both acinar and duct cells at 3 hours following IR (Fig. S3A, B), as previously reported (Meyer et al., 2017). At 24- and 48 hours post-IR, γH2AX foci were predominantly localized to duct cells (Fig. S3C, D), in agreement with previous reports (Marmary et al., 2016; Varghese et al., 2018). Irradiation using SARRP also induced high numbers of γH2AX foci by 3 hours, which was decreased in IR SMGs after 48 hours (Fig. S3E-H). These data were quantified and confirm that γH2AX foci were more prevalent in duct than in acinar cells at 48 hours following IR, indicating an elevated level of DNA damage in this cell population (Fig. S3I). Consistent with their DNA repair activity, mRNA expression levels of *Tgf-b1, Foxo3a*, and *Gadd45a* were rapidly, but transiently, increased by 3 hours post-IR (Fig. S4A-C).

The expression level of pro-inflammatory cytokines was assessed in SMG irradiated with the Cs source, using quantitative real-time PCR (qPCR). Expression of interleukin-1β (*Il-1β*) and tumor necrosis factor-α (*Tnf-α*) mRNAs were significantly increased at 3 hours post-IR (Fig. 3A, B). Expression of *Cxcl-2* mRNA, a pro-inflammatory cytokine produced by macrophages and demonstrated to be up-regulated in oral tissues after radiation (Shen et al., 2018), was also significantly increased at 3 and 24 hours post-IR (Fig. 3C). *P2y*_*2*_*R*, a member of the purinergic receptor gene family, shown to be up-regulated by Il-1β in salivary glands after injury (Turner et al., 1997; Khalafalla et al., 2020), was also increased (data not shown).

**Fig. 3.**
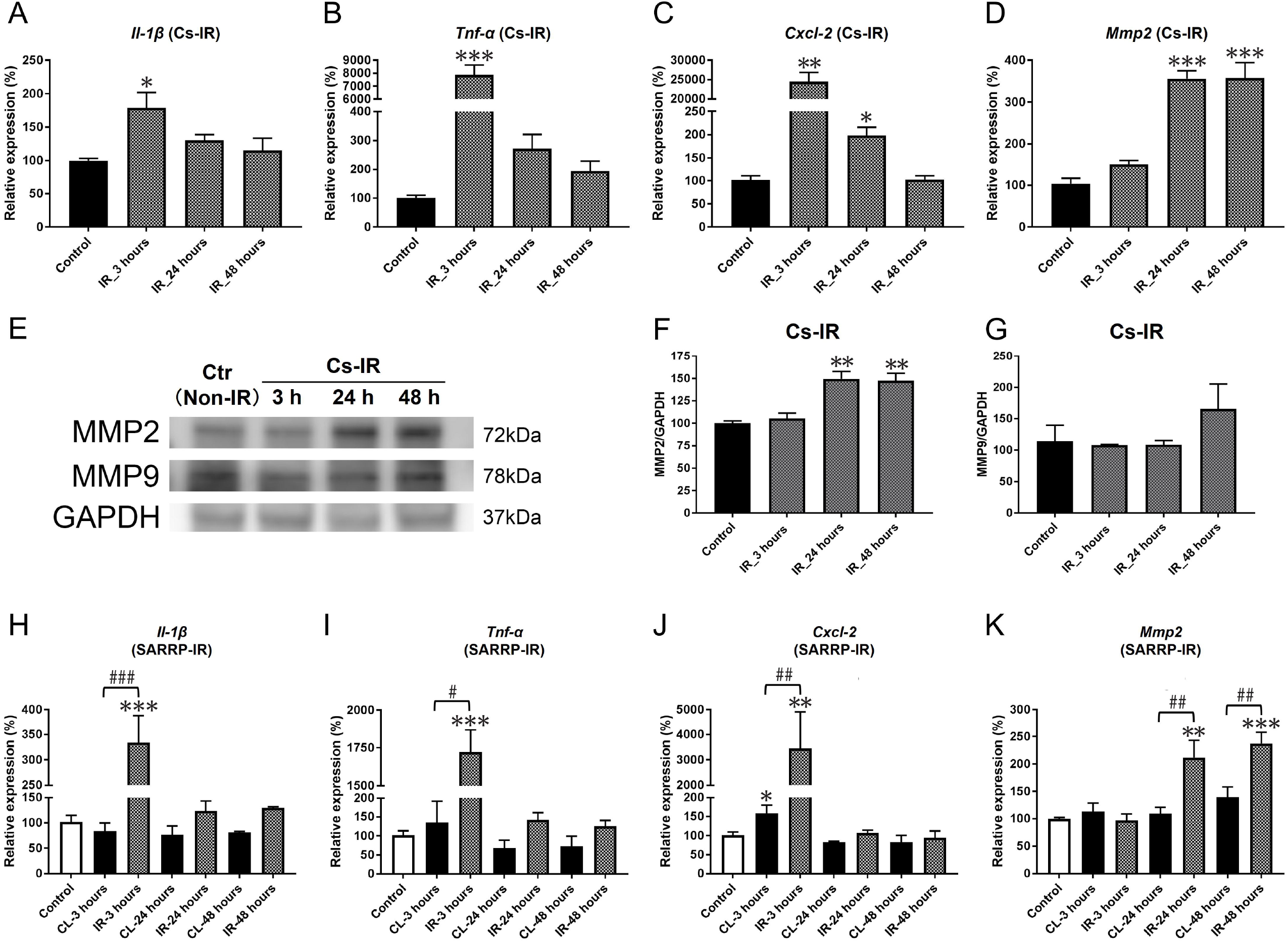
DNA damage and inflammatory factors are rapidly up-regulated following irradiation. **A**,**B**, Expression of *Il-1β* and *Tnf-α* mRNAs was up-regulated by 3 hours post-IR using the Cs source (* p = 0.026, ***p < 0.001, respectively). At 24 hours, expression of both cytokines had decreased compared to non-IR control. **C**, *Cxcl-2* mRNA was significantly increased at 3 and 24 hours post-IR (**p = 0.002, *p = 0.013), but decreased to control levels by 48 hours post-IR. **D**, Expression of *Mmp2* mRNA was increased at 24 and 48 hours post-IR (***p < 0.001), compared to non-IR control. **E**, Western blot analysis of MMP2, MMP9 and GAPDH protein abundance in control and irradiated SMGs (n = 3). **F**, MMP2 protein was increased at 24 and 48 hours post-IR (**p = 0.002). **G**, In contrast, MMP9 protein level did not change post-IR (p > 0.1). **H-K**, Following SARRP irradiation, expression profiles for (**H**) *Il-1β* (*** p < 0.001 vs. control), (**I**) *Tnf-α* (*** p < 0.001 vs. control), (**J**) *Cxcl-2* (** p = 0.015 vs. control) and (**K**) *Mmp2* (** p = 0.002, ***p < 0.001 vs. control) were similar to those induced using the Cs source. Error bars indicate mean ± SEM. Statistical analysis compared to control was performed using one-way ANOVA with Dunnett’s post-hoc test. Two-way ANOVA with Bonferroni test was used to compare IR and CL SMG (n = 3-5 each group): #p < 0.05, ##p < 0.01, ###p < 0.001.

Matrix metalloproteases, such as Mmp-2 and Mmp-9, are involved in extracellular matrix tissue remodeling, a process linked to inflammation (Duarte et al., 2015), and are stimulated by IR in some cell types (Wang et al., 2000; Lombaert et al., 2020). In bilaterally irradiated SMG, *Mmp2* mRNA expression was significantly increased at 24 and 48 hours post-IR (Fig. 3D), and MMP2 protein abundance was increased on western blots, relative to control protein (Fig. 3E, F). In contrast, there was no change in expression of *Mmp9* mRNA (data not shown) or protein abundance following IR (Fig. 3E, G). The upregulation of MMP2 may also be linked to the edema-like morphological changes observed in the irradiated SMGs (Fig. S2C, D).

Similar to bilaterally irradiated SMG, increased *Il-1β* and *Tnf-α* mRNA levels were observed after unilateral IR using the SARRP (Fig. 3H, I). *Cxcl-2* mRNA was also elevated at 3 hours post-IR in comparison to controls (Fig. 3J). *Mmp2* mRNA expression was increased significantly at 24 and 48 hours (Fig. 3K). Notably, although both radiation sources elicited similar pro-inflammatory responses, expression of these genes was not significantly altered in the CL SMGs (Fig. 3H-K).

### Irradiation transiently disrupts expression of mitochondrial factors

Since IR rapidly disrupts mitochondrial function in SMGs (Liu et al., 2017; Kawamura et al., 2018), we used qPCR to look for changes in the expression of key factors involved in mitochondrial biogenesis or ROS regulation. *Sirt3*, which plays a central role in maintaining mitochondrial homeostasis after stress (Marcus and Andrabi, 2018), was transiently down-regulated after IR, whereas Sirt1 and Sirt7, which are localized to the nucleus and linked to DNA repair (Finkel et al., 2009; Vazquez et al., 2016), were not altered (Fig. 4A; Fig. S4D, E).

**Fig. 4.**
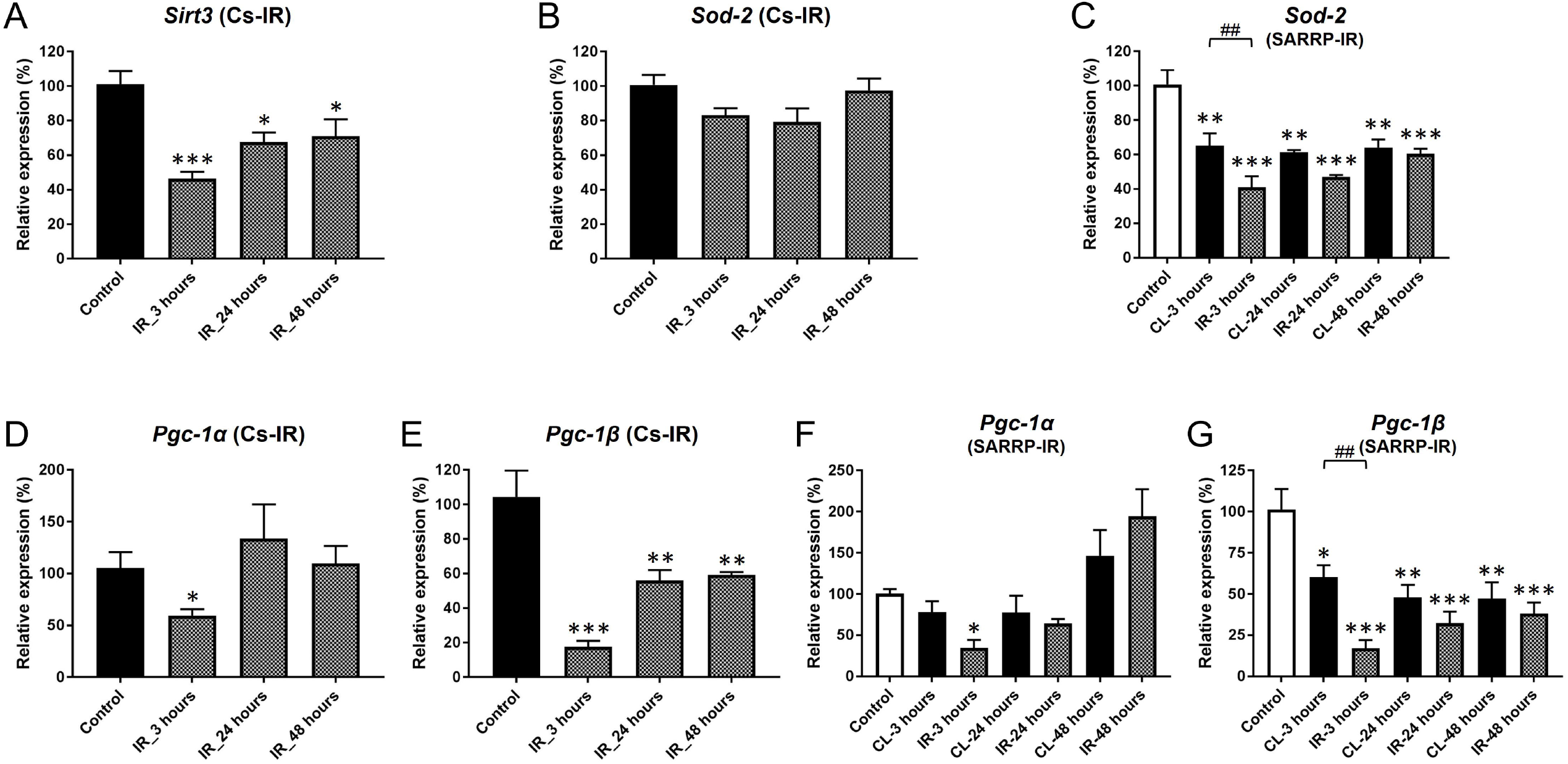
Irradiation transiently disrupts expression of mitochondrial factors. **A**, Expression of *Sirt3* was rapidly decreased by 3 hours after IR with the Cs source (***p < 0.001, *p < 0.025) ]), compared to the non-IR control. **B**, *Sod-2* mRNA levels did not significantly change after IR using the Cs source (one-way ANOVA; F = 2.822, p = 0.072). **C**, In contrast, *Sod-2* mRNA expression was significantly decreased after IR using SARRP, in both IR and CL SMGs by 3 hours vs. control (** p = 0.001, *** p < 0.001 vs. control), with lower expression in the IR-treated SMG (##p < 0.01). **D**, Expression levels of *Pgc-1*α mRNA was decreased by 3 hours (p = 0.037), but recovered by 24 hours (p = 0.997 [24 hours], p = 0.998 [48 hours]). **E**, *Pgc-1β* mRNA expression was significantly decreased by 3 hours (*p < 0.001) and remained at a lower level than the non-IR control up to 48 hours. **F**, A similar expression profile for *Pgc-1*α mRNA (* p = 0.027) was induced using SARRP. **G**, IR using SARRP significantly reduced expression of *Pgc-1β* in both IR (*** p < 0.001 vs. control) and CL SMGs (* p = 0.012, ** p = 0.002), with significantly lower expression in the IR-treated SMG (##p < 0.01 vs. CL). Error bars indicate mean ± SEM. Statistical analysis was performed compared to control (non-IR) using one-way ANOVA with Dunnett’s post-hoc test, or comparing IR and CL SMG using two-way ANOVA with Bonferroni test (n = 3-5 each group).

Expression of *Sod-2* mRNA, a mitochondrial superoxide dismutase that counters oxidative stress (Wang et al., 2018), did not change within 48 hours after IR using the Cs source (Fig. 4B), but was significantly decreased in both SARRP-irradiated and CL SMGs (Fig. 4C). Expression of *Sod-1*, a cytoplasmic protein, was not altered by IR administered from either source (Fig. S4F; and data not shown). The transcriptional coactivators PGC-1 alpha and PGC-1 beta are key regulators of mitochondrial biogenesis, and were both down-regulated within 3 hours after IR (Fig. 4D, E). mRNA expression of *Pgc-1a* and *Pgc-1b* was also decreased at 3 hours in SARRP-irradiated SMGs (Fig. 4F, G). Notably, *Pgc-1b* expression was also significantly decreased in CL SMGs compared to control glands (Fig. 4G). Thus, IR both directly and indirectly disrupted expression of several mitochondrial factors.

### Pro- and anti-apoptotic gene expression is induced by radiation

Irradiation of the salivary glands leads to activation of p53 (Limesand et al., 2006), which can result in cell cycle arrest or apoptosis (Riley et al., 2008). A previous study reported that the p53-dependent pro-apoptotic factors *Bax* and *PUMA* were upregulated at 4 and 8 hours post-IR (Avila et al., 2009). Consistent with these observations, our data showed that *Bax* mRNA was significantly up-regulated at 3, 24 and 48 hours after IR using the Cs source (Fig. 5A). However, histological analysis of irradiated rat and mouse SMGs within 48 hours post-IR did not reveal evidence of apoptosis (Choi et al., 2009; Marmary et al., 2016). We investigated whether additional targets of p53, including the anti-apoptotic factors *Bcl-2, Bcl-xl* and *p21*, were upregulated. mRNA expression levels of *Bcl-2* and *Bcl-xl* showed transient elevation by 3 hours post-IR, but returned to control levels by 24 hours (Fig. 5B, C). Expression of p21 was rapidly and significantly up-regulated by 3 hours post-IR, and remained high through 48 hours (Fig. 5D). The expression profiles of these genes were similar after IR using SARRP (Fig. 5E-H). Notably, with the exception of p21, there was not a significant effect of IR on these genes in the CL SMGs.

**Fig. 5.**
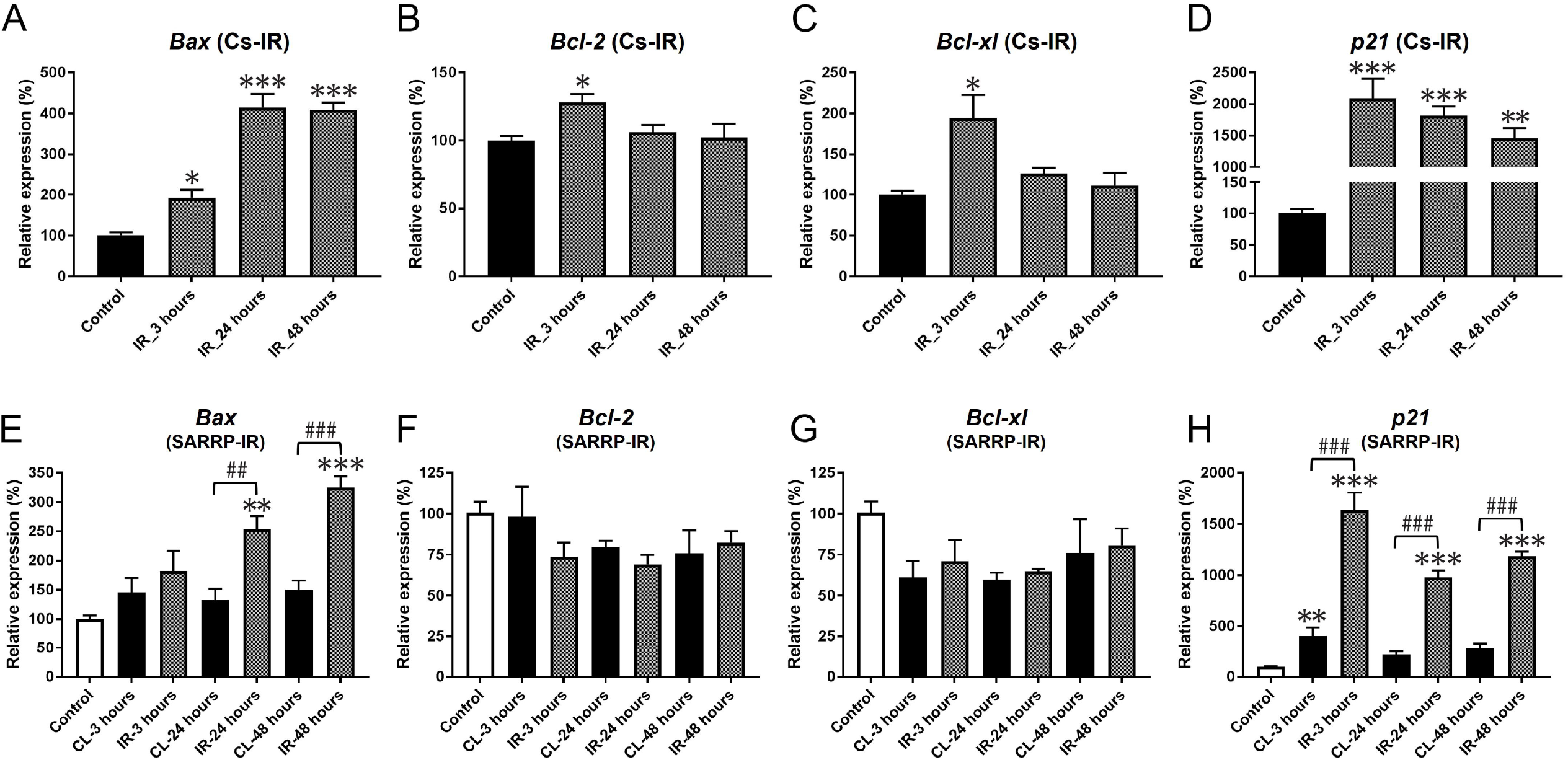
Pro- and anti-apoptotic gene expression is induced by radiation. **A**, *Bax* mRNA expression was significantly elevated at 3 hours (* p = 0.022), 24 hours and 48 hours (*** p < 0.001) after IR using the Cs source compared to the non-IR control. **B**,**C**, Anti-apoptotic factors Bcl-2 and Bcl-xl were transiently elevated by 3 hours (* p = 0.024; * p = 0.003, respectively). **D**, Expression of *p21* mRNA was significantly increased by 3 hours (*** p < 0.001), and remained elevated at 24, and 48 hours (** p = 0.001). **E-H**, After IR using SARRP, expression profiles of Bax, Bcl-2, Bcl-xl and p21 were similar to those observed using the Cs source. **E**, Bax mRNA levels were elevated at 24 (** p = 0.002) and 48 (***p < 0.001) hours, and were significantly higher in IR-treated SMG compared to CL SMG (##p < 0.01, ###p < 0.001). **F**,**G**, Expression levels of *Bcl-2* and *Bcl-xl* mRNA did not change significantly following IR using SARRP. **H**, *p21* mRNA expression was significantly elevated at 3 hours in IR-treated and CL SMGs (**p = 0.005, *** p < 0.001 vs. control), and expression remained elevated in the IR-treated SMG up to 48 hours (*** p < 0.001 vs. control). *p21* mRNA levels in the IR-treated SMG were significantly higher than in the CL SMG (###p < 0.001). Error bars indicate mean ± SEM. Statistical analysis was performed compared to control (non-IR) using one-way ANOVA with Dunnett’s post-hoc test, and to compare IR and CL SMG using two-way ANOVA with Bonferroni test. (n = 3 - 5 each group).

## Discussion

Radiation-induced effects in mouse salivary glands are divided into short- and long-term changes (Coppes et al., 2001; Jasmer et al. 2020). This study was undertaken to search for rapid changes in gene or protein expression that may yield insights into IR-induced causes of hyposalivation. Murine SMGs were irradiated bilaterally using a ^137^Cs gamma ray irradiator, or unilaterally, using the SARRP, an image-guided microirradiator. Radiation from the cesium source is in the form of gamma rays (with an energy of 662 keV), while the SARRP delivers 225 kVp X-rays with the beam filtered to remove low-energy photons. Both are categorized as low linear energy transfer (LET) ionizing irradiation, and result in similar, although not always identical, physiological effects (Iyer and Lehnert, 2000; Iyer and Lehnert, 2002; International Agency for Research on Cancer, 2000). Our data show that IR-induced changes are similar with either low-LET gamma or X-rays. Moreover, we report radiation-induced coordinated bystander effects in non-irradiated SMGs.

Acute hyposalivation in mice is observed by 3 days after IR (Avila et al., 2009; Grundmann et al., 2010; Morgan-Bathke et al., 2014). We measured a significant decrease in saliva volume at 3 hours following bilateral IR, and found that secretion was recovered at 24 hours. At 48 hours, secretion was again decreased, and after a modest recovery at 1-2 weeks, continued to decline.

Increased extracellular space and fluid accumulation seen in SMGs after IR suggest disruption of the epithelial barrier. Alterations in expression of the tight junction genes Claudin-3 and -4 occurs within 3 days of IR in parotid glands (Yokoyama et al., 2017), and increased Claudin-4 expression accompanies acute lung injury (Wray et al., 2009). We found that *Claudin-3* and *Claudin-4* mRNAs were up-regulated as early as 3 hours post-IR, which should preserve intracellular junctions. However, claudins promote the activation of pro-MMP2 (Miyamori et al., 2001; Lee et al., 2008; Hwang et al., 2010), which degrades type IV collagen, a major component of the basement membrane (Araya et al., 2001; Zhao et al., 2004). MMP2 was increased in minipig parotid glands (Lombaert et al., 2020), and in IR-treated SMGs at 24 hours, consistent with increased intercellular edema.

Radiation induces the activation of p53 and the pro-apoptotic factors, Bax and Puma, by 4 hours post-IR (Limesand et al., 2006; Avila et al., 2009). BAX homodimers accelerate apoptosis by inducing release of cytochrome c; whereas, heterodimers of BAX with the anti-apoptotic proteins Bcl-2 or Bcl-xl, prevent caspase activation (Reed, 1994; Kim, 2005). By 3 hours post-IR, Bcl-2 and Bcl-xl are both upregulated, as is p21, which functions to suppress apoptosis, in part by inhibiting the activity of caspases (Sohn et al., 2006; Mirzayans et al., 2013). The transient up-regulation of Bcl-2 and Bcl-xl, together with p21, may limit apoptosis, which declines within 48 hours post-IR (Avila et al., 2009).

IR induced a transient decrease in the transcription factor Mist1, which maintains the secretory phenotype of acinar cells (Pin et al., 2001; Lo et al., 2017). Mist1 regulates the expression of *Aqp5*, a water channel critical for saliva secretion (Jia et al., 2008), and *Cx32*, a gap junction protein in exocrine acinar cells (Rukstalis et al., 2003). The sensitivity of the MIST1 transcription factor to stress or injury (Karki et al., 2015) most likely accounts for the down-regulation of all three mRNAs at 3 hours post-IR. Consistent with earlier reports in rat SMG (Takagi et al., 2003; Li et al., 2006), AQP5 protein levels decreased within 48 hours.

IR has been linked to disruptions in calcium signaling and mitochondrial pathways in acinar cells (Liu et al., 2017). Sirtuins are histone deacetylases, which influence cellular responses to external signals by regulating cell cycle, metabolism and genome stability (Chalkiadaki and Guarente, 2015). A previous study found that Sirt1 mRNA was significantly increased in mouse parotid glands within 30 minutes after IR with 5 Gy (Meyer et al., 2017). However, there was no significant change in Sirt1 or Sirt7 mRNA levels in SMG at 3 hours after 15 Gy IR. The discrepancy may be due to different radiation doses, time of analysis, or salivary gland type. In contrast, Sirt3, a mitochondrial protein that coordinates mitochondrial metabolism (Giralt and Villarroya, 2012), thereby limiting levels of ROS (Chalkiadaki and Guarente, 2015), was transiently down-regulated at 3 hours post-IR. The expression of SOD genes involved in ROS homeostasis (Zelko et al., 2002) was not changed, but the coactivators PGC-1α and PGC-1β, which regulate the SOD genes (Lin et al., 2005), were down-regulated by 3 hours post-IR. Further investigation into gene expression changes within these rapidly responsive pathways is warranted.

ROS produced by low LET irradiation can migrate to distant sites through cell contacts, or across cell membranes, causing responses in non-targeted cells (Desouky et al., 2015). The precision of SARRP allowed us to investigate off-target effects induced in the CL SMG. Unilateral IR resulted in gene expression changes not only in the irradiated, but also in the CL SMG. Similar off-target effects were observed in the non-irradiated parotid glands of minipigs (Lombaert et al., 2020). In addition to ROS, release of soluble factors including nitric oxide and activated cytokines from irradiated cells also contributes to bystander effects (Prise and O’Sullivan, 2009; Najafi et al., 2014). We speculate that the elevated expression of the cytokines IL-1b, TNF-α and Cxcl2 by 3 hours after IR is involved in the propagation of bystander effects to the CL SMG. Although not investigated in this study, purinergic receptors are also likely to play a role in promoting bystander effects (Jasmer et al. 2020).

Importantly, our results demonstrate that non-irradiated contralateral SMGs undergo the same responses as irradiated glands following unilateral IR. Many of the changes in gene expression detected in the irradiated SMG, including *Mist1, Aqp5, Pgc-1β* and *Sod-2*, were also observed in the non-irradiated CL SMG. Although the coupling mechanism is not understood, it is known that unilateral injury or stress results in a similar response in both SMGs (Walker and Gobé, 1987; Lombaert et al., 2020). An understanding of how radiation impacts salivary gland function must include the response of non-irradiated glands to bystander effects. Further investigation into the mechanisms involved is required for rational design of radioprotective strategies.

## Methods and Materials

### Animals

Female C57BL/6J (Jackson Laboratory, ME, USA) mice aged 4--12 weeks old were used in this study. Animals were housed in groups and maintained in a 12 hour light/dark cycle with food and water available *ad libitum*. All procedures were approved and conducted in accordance with the University Committee on Animal Resources at the University of Rochester Medical Center.

### Irradiation

Two radiation sources were utilized to investigate the short-term effects of radiation. The ^137^Cs gamma (Cs) radiation source (Shepard), in combination with a custom-built brain-slit collimator of 4 mm, was used to deliver radiation bilaterally but to limit radiation exposure to the neck region of C57BL/6 female mice (Fig. S1A), as previously described (Weng et al. Cell Rep 2018). Mice were anesthetized with ketamine (90 mg/kg) and xylazine (9 mg/kg) via intraperitoneal injection and irradiated with a single dose of 15.0 Gy within 20 minutes. Control mice were administered ketamine/xylazine, but not irradiated.

The Small Animal Radiation Research Platform (SARRP) (Xstrahl) permits unilateral IR of a single SMG (Fig. S1B,C), through spatial targeting of the radiation area following a computed tomography (CT) scan (Fig. S1D-F). Mice were anesthetized with vaporized isoflurane through a nose cone, and a single dose of 15.0 Gy was targeted to the left SMG of irradiated mice, using a 10×10 mm collimator, as previously described (Bachman et al., 2020).

Mice irradiated with the Cs source were analyzed for saliva secretion, histology, and gene expression at shortly after IR. Mice irradiated using the SARRP were analyzed for histology, and gene expression to investigate bystander effects post-IR.

### Saliva collection

Total saliva was collected both before and after IR from the same mice. To establish the baseline of saliva volume, mice were anesthetized with ketamine (75 mg/kg) and xylazine (7.5 mg/kg) and their saliva secretion was stimulated with an intraperitoneal injection of the muscarinic receptor antagonist pilocarpine (0.5 mg/kg). Total saliva was collected for 20 minutes into pre-weighed tubes using glass capillary tubes placed into the oral cavity under the tongue. Total saliva volume was normalized to individual body weight (μL/g) at each time-point.

### Histological analysis

SMGs were dissected from control (non-IR), Cs-irradiated and SARRP-irradiated (both IR and CL SMGs) mice at 3, 24 and 48 hours post-IR and fixed in 4% paraformaldehyde at 4°C overnight. Mice used for saliva collection were not used for tissue collection. Staining was processed as previously described (Varghese et al., 2018; Weng et al., 2018). Briefly, fixed tissues were embedded in paraffin and sections were cut to 5 μm and stained with hematoxylin and eosin (H&E) for morphological assessment. For immunohistochemistry, antigen retrieval was performed in HIER buffer (10mM Tris-base, 1mM EDTA, pH 9.4 or citrate, pH 6.0). CAS-Block™ Histochemical Reagent (Thermo Fisher Scientific, 008120) was used to block for 1 hour. Primary antibodies: Aquaporin5 (AQP5, Abcam, 1:100), Claudin-3 (CLN3, Abcam, 1:500), Claudin-4 (CLN4, Abcam, 1:500), γH2AX (Millipore Sigma, 1:200), MIST1 (Abcam, 1:200), Muscarinic receptor type 3 (M3R, 1:200), Nkcc1 (Cell Signaling, 1:1000), ZO-1 (Thermo Fisher Scientific, 1:1000) and loading control: β-actin and GAPDH were applied overnight at 4°C. Secondary antibodies were donkey-anti-rabbit (Thermo Fisher Scientific, 1:1000) or donkey-anti-mouse (Thermo Fisher Scientific, 1:1000). Nuclei were stained with DAPI (D1306, Thermo Fisher Scientific).

Imaging and analysis were performed as previously described (Weng et al., 2018; Ingalls et al., 2020). Images for H&E staining were acquired using an Olympus DX41 microscope with a DP41 camera and analyzed using ImageJ software (National Institutes of Health, USA). Fluorescent images were acquired using Olympus IX85 phase contrast microscope at 40× magnification or a Leica TCS SP5 confocal microscope with 40× oil immersion objective and Argon laser. Fluorescent images were converted to 8-bit format, thresholded to binary, and a watershed function was used to distinguish individual cells. The number of nuclei (DAPI, NKCC1-positive and NKCC1-negative) and dots (γH2AX) were counted automatically using the particle analysis plugin.

### Total RNA extraction and quantitative real-time PCR

SMGs were dissected from control (non-IR), Cs-irradiated and SARRP-irradiated (both IR and CL SMGs) mice at 3, 24 and 48 hours post-IR. SMGs were dissected into TRIzol reagent and stored at -80°C until total RNA extraction. Total RNA was extracted using the OMEGA kit (Omega Bio-tek) and reverse-transcribed using the iScript™ cDNA synthesis kit (Bio-Rad), according to the manufacturer’s instructions. Quantitative real-time PCR (qPCR) analysis of individual cDNAs was processed on a CFX96TM Real-Time System (Bio-Rad) using SsoAdvanced™ Universal SYBR Green Supermix (Bio-Rad) and the following PCR primer sets: mouse *Rps29* (reference gene), *Mist1, Aqp5, M*_*3*_*r, P2y2, Il-1β, Tnf-α, Cxcl-2, Mmp2, Bax, Bcl-2, Bcl-xl, p21, Sirt1, Sirt3, Sirt7, Foxo3a, Gadd45a, Tgf-*β*1, Sod-1, Sod-2, Pgc-1*α and *Pgc-1β* (primer sequences in Table 1). Target genes were normalized to mouse *Rps29* as a reference gene. Reference and target genes were only compared from the same plate. QPCR results were analyzed by the 2^-*ΔΔ*Ct^ method. C_t_ values of less than 35 were obtained from all target genes. All genes were measured from n = 3-5 mice. cDNA samples were tested using biological duplicates (2 wells per primer set per sample). A series experiment (control, 3, 24 and 48 hours post-IR using Cs-source, or control [non-IR], CL and IR-SMGs at 3, 24 and 48 hours post-IR using SARRP) was run using several primer sets.

**Table 1.**
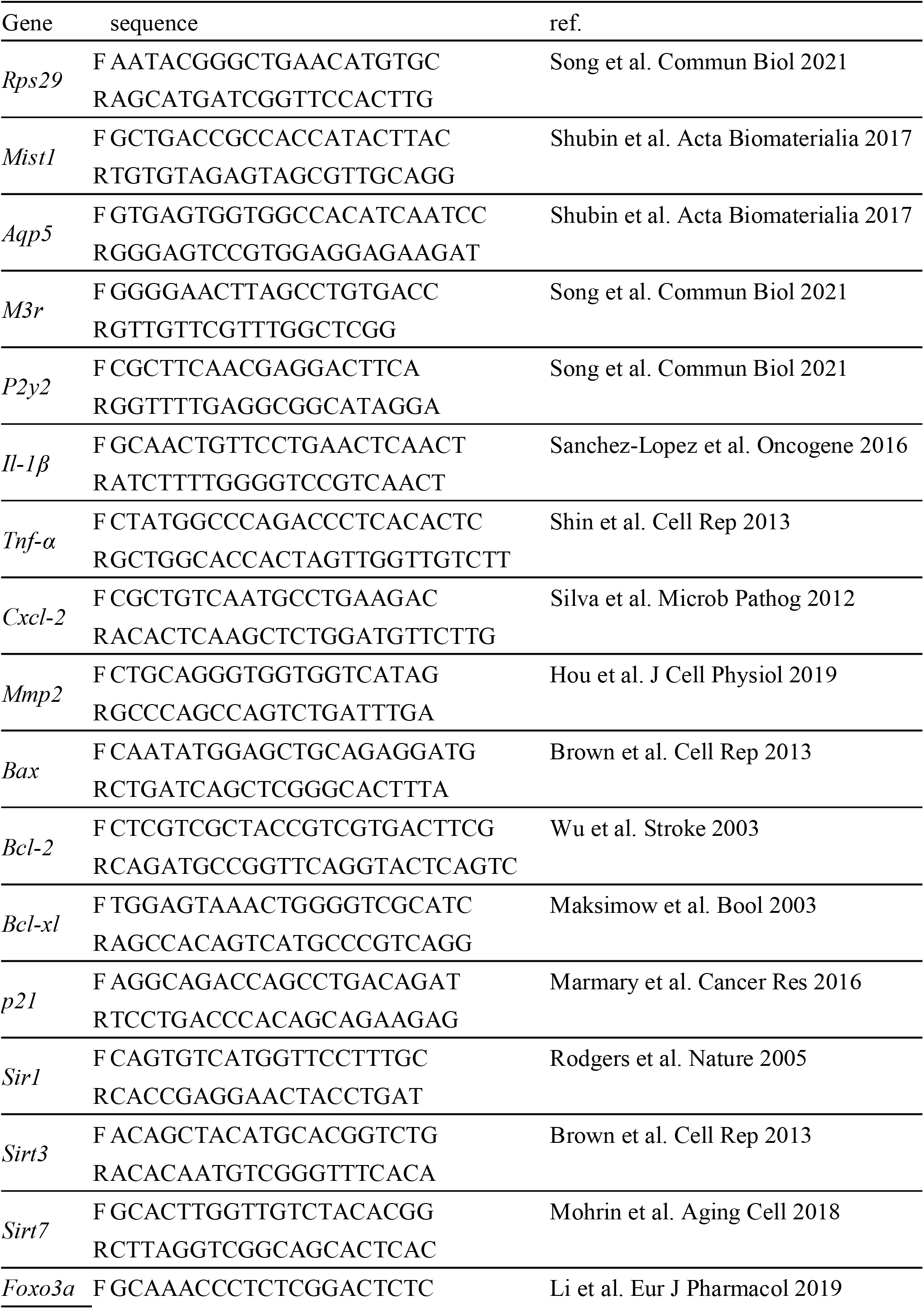
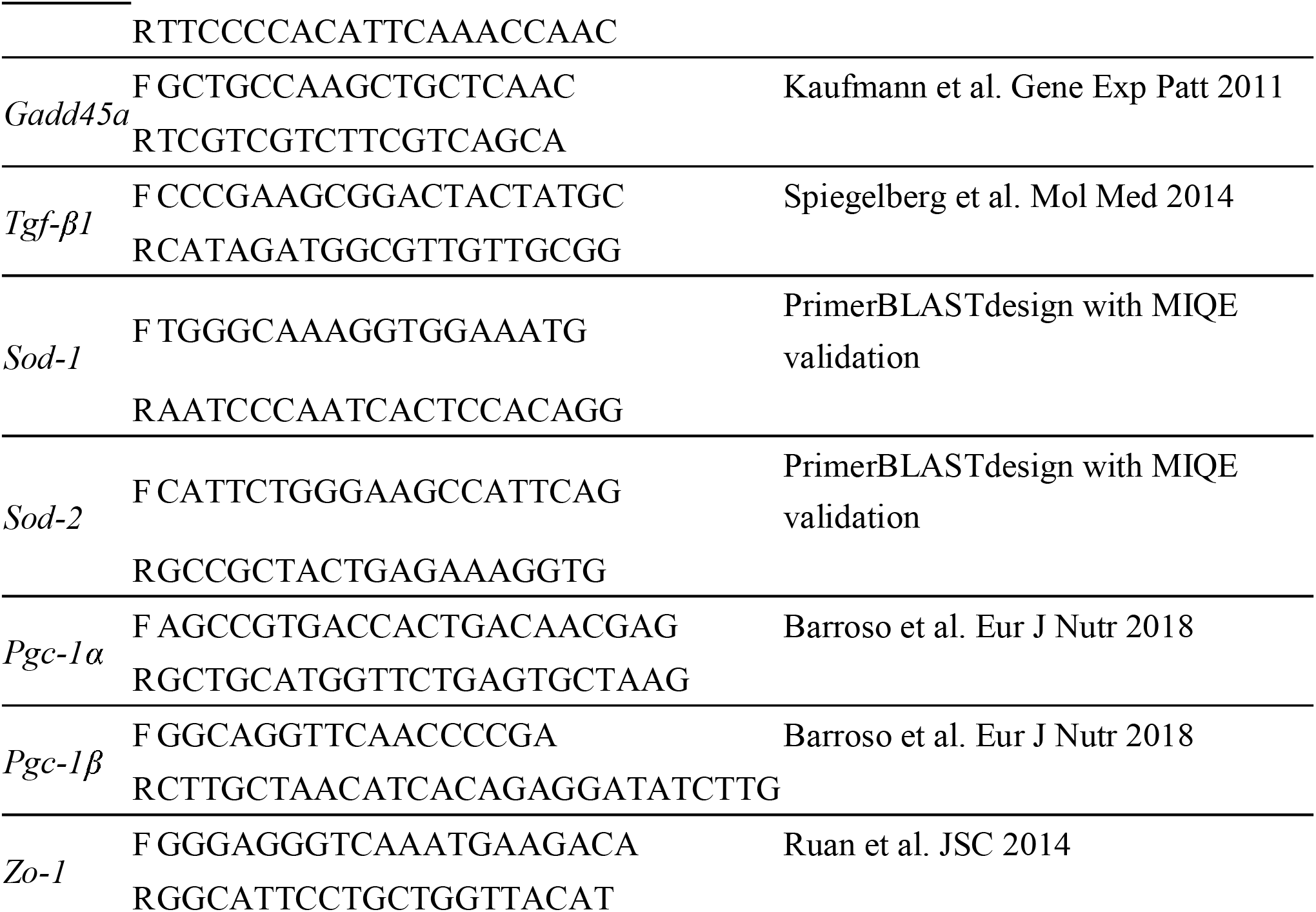
Primer sequences for qPCR.

### Western blot analysis

Total protein was extracted from the SMGs of control (non-IR) and Cs-irradiated mice at 3, 24, and 24 hours post-IR as previously described (Li et al., 2006). Briefly, isolated SMGs were placed in chilled protein extraction buffer (50 mM Tris-HCl [pH 8.0], 2 mM EDTA, 250 mM sucrose, 1 mM β-mercaptoethanol and protease/phosphatase inhibitor (Cell Signaling Technology, MA, USA). The tissue was homogenized for 1 min on ice. The homogenate was centrifuged at 2,000 rpm for 15 min at 4°C and the supernatant was collected as the total protein fraction. The protein concentration of each sample was measured at 750 nm using the Bio-Rad reagent according to the manufacturer’s protocol. 20 µg of total protein was run on a MINI-PROTEIN TGX pre-cast gel (Bio-Rad, 4561093) and followed by protein transfer onto PVDF membrane (Bio-Rad, 1620174). The membrane was blocked in 2% BSA-TBST or 5% skim milk-TBST at room temperature for 1 hour. The following primary antibodies were applied overnight at 4°C: AQP5 (Abcam, 1:1000), CLN3 (Thermo Fisher Scientific, 1:1000), CLN4 (Abcam, 1:1000), Connexin 32 (CX32, Thermo Fisher Scientific, 1:1000), E-CAD (Biosciences, 1:3000), MIST1 (Abcam, 1:1000), Matrix Metalloprotease 2 (MMP2, Thermo Fisher Scientific, 1:1000), Matrix Metalloprotease 9 (MMP9, Thermo Fisher Scientific, 1:1000), β-actin (Santa Cruz, 1:3000) and GAPDH (Thermo Fisher Scientific, 3 µg/mL). Appropriate secondary antibodies (an anti-rabbit IgG [MP-7401, Vector] or an anti-Mouse IgG [1706516, Bio-Rad]) were performed to detect the proteins. The signals were detected using Pierce™ ECL Western Blotting Substrate (32106, Thermo Fisher Scientific). For the re-probing of loading control (β-actin or GAPDH), the stained membrane was washed with TBST and incubated in Restore Western Blot Stripping Buffer (Thermo Fisher Scientific, cat #. 21059) for 30 min at room temperature. After 3 washes with TBST, the membrane was blocked in 2% BSA-TBST or 5% skim milk-TBST at room temperature for 1 hour. The primary antibody (β-actin or GAPDH) was applied overnight at 4°C. The membrane was incubated with anti-Mouse IgG [1706516, Bio-Rad] secondary antibody and the protein was detected using Pierce™ ECL Western Blotting Substrate (32106, Thermo Fisher Scientific). Protein levels were quantified and compared using ImageJ (National Institutes of Health, USA) densitometric analysis (Schneider et al., 2012).

### Statistical analysis

All results are presented as mean ± SD. Data for bilateral IR were analyzed by one-way analysis of variance (ANOVA) with Dunnett’s post-hoc test compared to control (non-IR) SMGs. Two-way ANOVA with Bonferroni post-hoc test was used to compare the CL and SARRP-irradiated SMGs using SPSS. Differences were considered significant at p < 0.05.

## Acknowledgements

We thank Dr. Brian Marples, Department of Radiation Oncology, for discussions and critical reading of the manuscript.

## Author contributions

Conceptualization: H.U., C.E.O; Methodology: H.U., M.H.I., E.M., C.J.J., E.H., C.E.O.; Validation: H.U.; Formal analysis: H.U.; Investigation: H.U., M.H.I., E.M.; Resources: H.U., M.H.I., E.M., C.J.J., E.H.; Data curation: H.U.; Writing-original draft preparation: H.U., C.E.O.; Writing-review and editing: H.U., M.H.I., E.M., C.J.J., E.H., R.C.F., C.E.O.; Visualization: H.U.; Supervision: C.E.O; Project administration: C.E.O.; Funding acquisition: H.U., M.H.I., C.E.O.

