## Supplemental Figure Legends for "Short-term and bystander effects of radiation on murine submandibular glands"

Uchida et al.

Supplemental Files

**Fig. S1. A,** At 3 months after IR, loss of fur color indicates the irradiated area. Irradiation using the Cs source with the brain-slit collimator delivers a dose of 15 Gy to the entire neck region, including both SMG. **B,C,** At 3 months after IR, mice irradiated unilaterally using the SARRP show loss of fur color within a defined region including only a single SMG. **D-F,** CT images show the area receiving radiation using SARRP. Blue lines delineate area receiving radiation dose of approximately 15 Gy. Red line and red dots indicate the isodose line and isocenter, respectively. Outside the area indicated, the radiation dose is close to 0 Gy.

**G,** Timeline for saliva collection. Open arrows mark collection time points. **H, I,** Base line saliva volume was determined in all mice prior to IR (open boxes). Saliva volume from non-IR mice (blue boxes) and from IR mice (yellow boxes) was collected at the timepoints shown. **H,** Saliva volume collected from non-IR mice was not changed compared to baseline saliva measured in all mice (one-way ANOVA; F = 1.246, p = 0.251, baseline: 1^st^ day; n = 12, 2^nd^ and 3^rd^ day; n = 13, non-IR, 3 hours; n = 3, 24 hours; n = 6, 48 hours; n = 3, 1^st^, 2^nd^, 4^th^, 5^th^, 7^th^ – 12^th^ weeks; n = 6, 3^rd^ weeks; n = 4, 6^th^ weeks; n = 5 mice). **I,** Saliva volume collected from IR mice decreased significantly in a time-dependent manner (one-way ANOVA; F = 14.296, p < 0.001, IR mice, 3 hours; n = 5, 24 hours; n = 6, 48 hours; n = 4, 1^st^ – 3^rd^, 5^th^ – 12^th^ weeks; n = 7, 4^th^ weeks; n = 6 mice). Data plotted using box-and-whisker plots. *p < 0.05, **p < 0.01, ***p < 0.001 (compared to baseline: one-way ANOVA with Dunnett’s post-hoc test).

**Fig. S2.** Hematoxylin and eosin (H&E) stain of control (**A**) and irradiated SMG at (**B**) 3 hours, (**C**) 24 hours and (**D**) 48 hours following IR using the Cs source. Outlined areas are shown below at higher magnification. Scale bars = 50 μm. H&E staining of SMG unilaterally irradiated using the SARRP at (**E)** 3 hours, (**F**) 24 hours and (**G**) 48 hours following IR. Outlined areas are shown below at higher magnification. H&E staining of contralateral (CL) SMG from mice irradiated using the SARRP at (**H**) 3 hours, (**I)** 24 hours, and (**J**) 48 hours after IR. Outlined areas are shown below at higher magnification. Scale bars = 50 μm.

**Fig. S3.** **A,** SMG were isolated from control or from irradiated mice at (**B**) 3 hours, (**C**) 24 hours and (**D**) 48 hours after IR using the Cs source and stained with γH2AX (green), and NKCC1 (red). Nuclei were stained with DAPI (blue). **E,** Contralateral (CL) and irradiated SMGs were isolated from mice at (**F**) 3 hours, (**G**) 24 hours, and (**H**) 48 hours after unilateral IR using SARRP and stained with antibody to γH2AX (red). Nuclei were stained with DAPI (blue). Scale bars = 50μm. **I,** Quantification of γH2AX foci in acinar (NKCC1-positive) (mean ± SD, p = 0.011 [3 hours], p = 0.564 [24 hours], p = 1.000 [48 hours] vs. control) and duct (NKCC1-negative) cells (mean ± SD, p < 0.001 [3 hours], p = 0.002 [24 hours], p = 0.007 [48 hours] vs. Control, p < 0.001 [24 hours], p < 0.001 [48 hours] vs. 3 hours post-IR) (n = 3 mice, two-way ANOVA; p = 0.765 [control], p < 0.001 [3 hours], p = 0.008 [24 hours], p = 0.012 [48 hours]). Statistical analysis compared to control was performed using one-way ANOVA with Dunnett’s post-hoc test, or to compare IR-treated to CL SMG using two-way ANOVA with Bonferroni test (n = 3 each group): *p < 0.05, **p < 0.01, ***p < 0.001 (one-way ANOVA), $$$ p < 0.001 compared to 3 hours post-IR or #p < 0.05, ##p < 0.01, ###p < 0.001 (two-way ANOVA).

**Fig. S4. A-C,** Expression levels of (**A**) *Tgf-b1 (*p = 0.040), (**B**) *Foxo3a* (p = 0.017) and (**C**) *Gadd45a (*p = 0.034) mRNAs were transiently increased by 3 hours post-IR but returned to background levels by 24 hours. **D,E,** *Sirt1* and *Sirt7* levels were not changed significantly following IR (*Sirt1*: p = 0.831 [3 hours], p = 0.082 [24 hours], p = 0.043 [48 hours], *Sirt7*: p = 0.549 [3 hours], p = 0.983 [24 hours], p = 0.983 [48 hours]). **F,** Similarly, expression of *Sod-1* mRNA, which encodes a cytoplasmic protein, was not changed within 48 hours post-IR (*Sod-1*, one-way ANOVA; F = 1.671, p = 0.213). Data show mean ± SEM. Statistical analysis was performed using one-way ANOVA with Dunnett’s post-hoc test (n = 3-5 each group): *p < 0.05, ***p < 0.001 compared to control (non-IR).
