## Supplementary figures and images for "Short-term and bystander effects of radiation on murine submandibular glands"

### Figure S1.

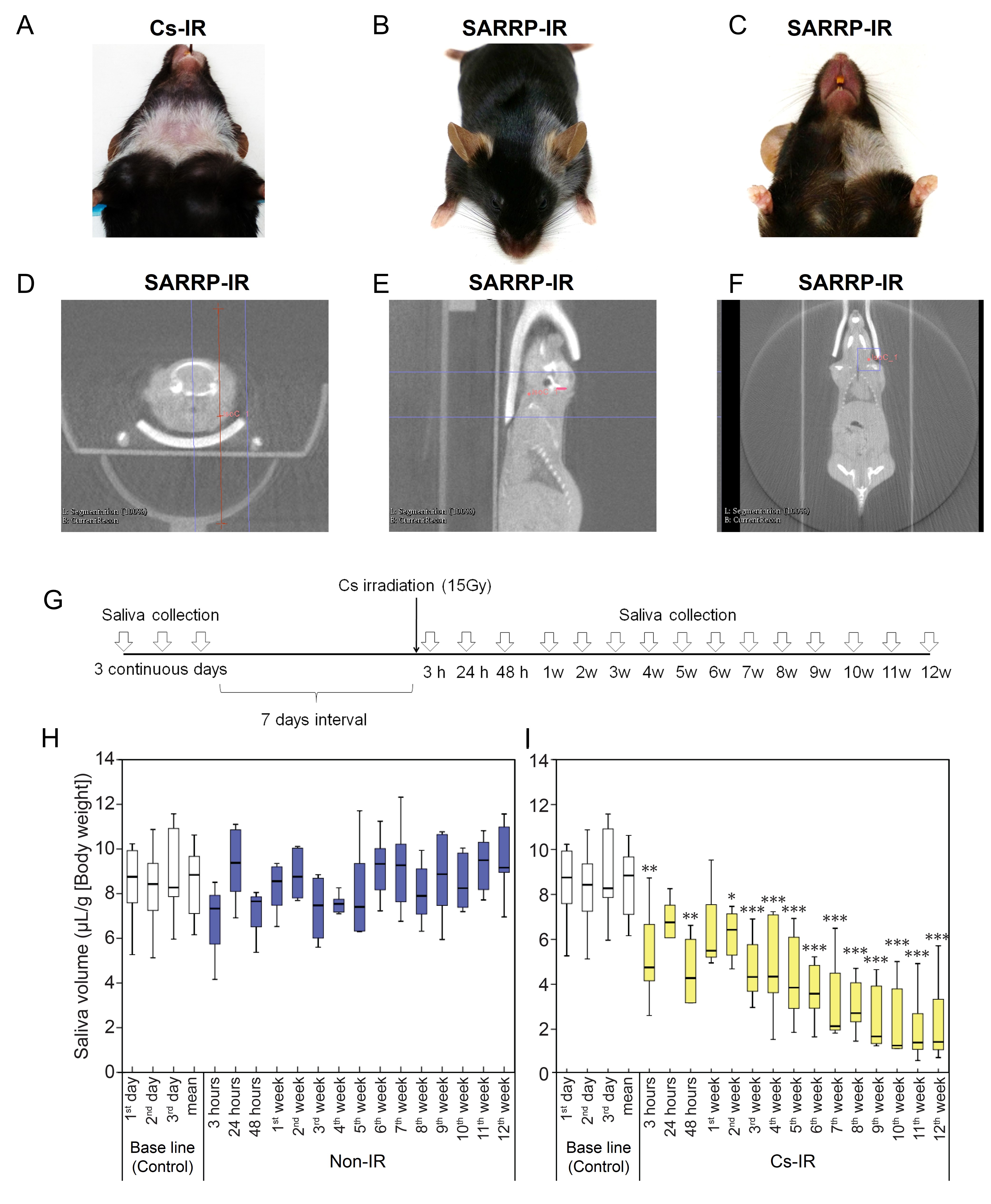

### Figure S2.

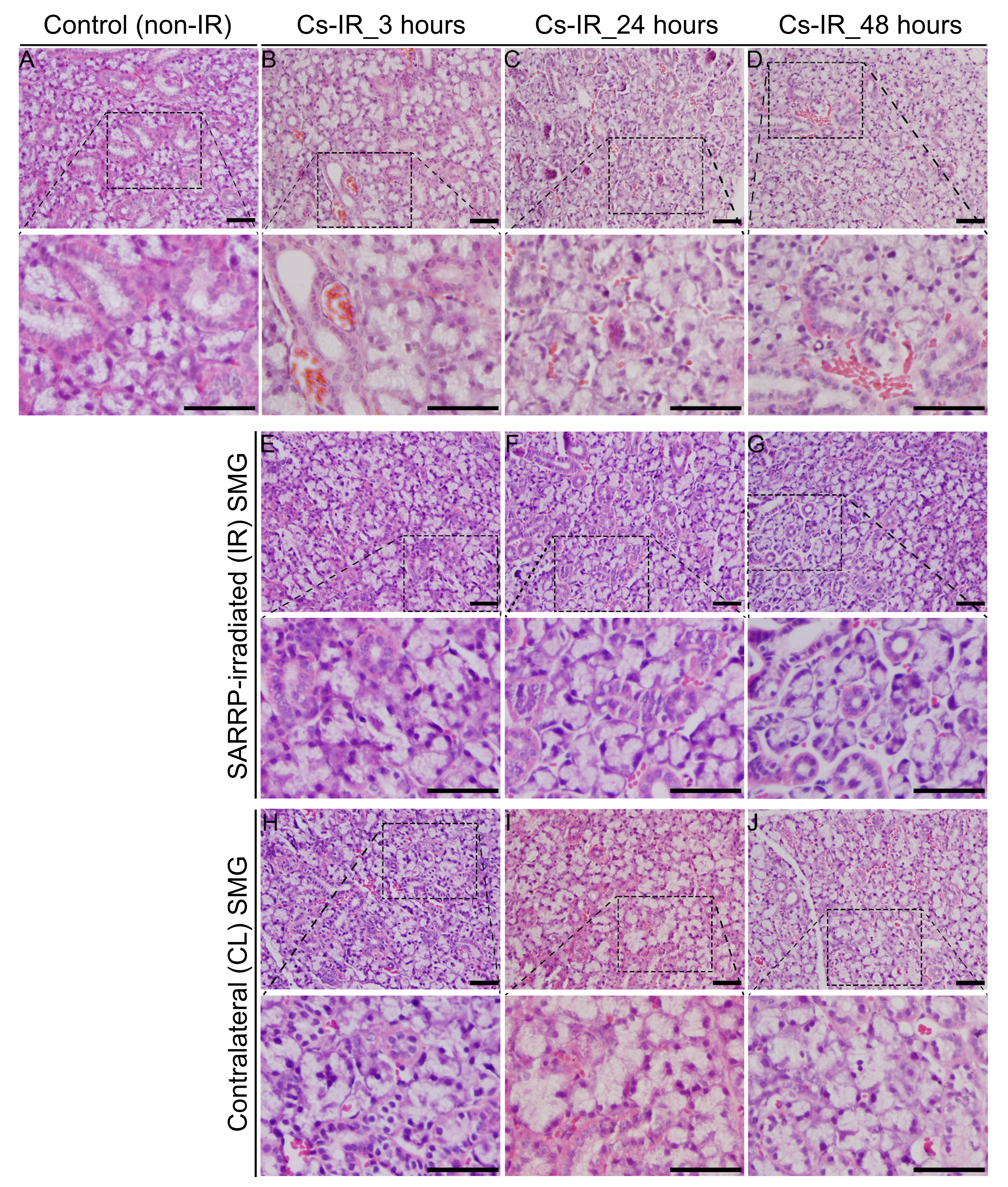

### Figure S3.

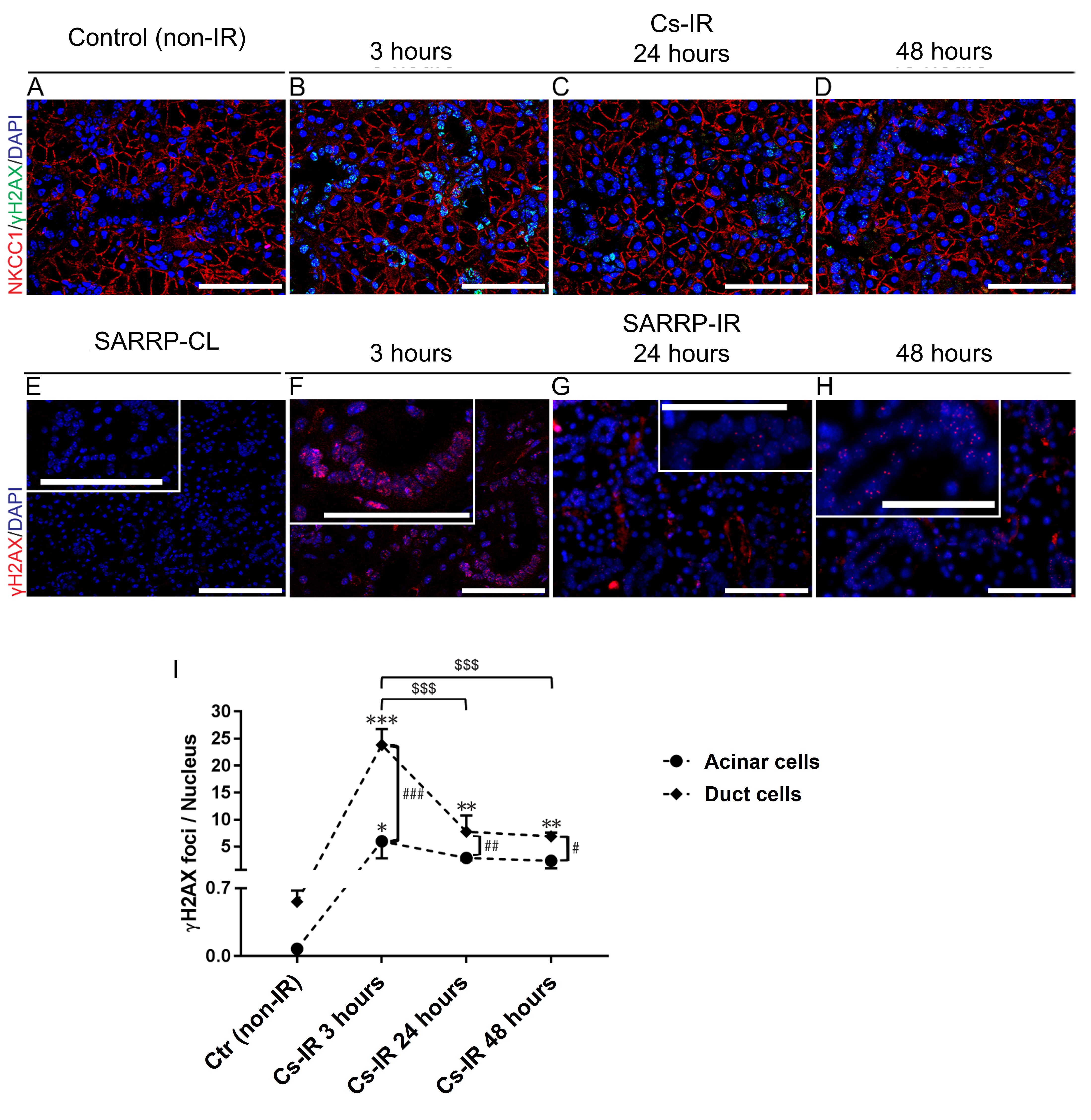

### Figure S4.

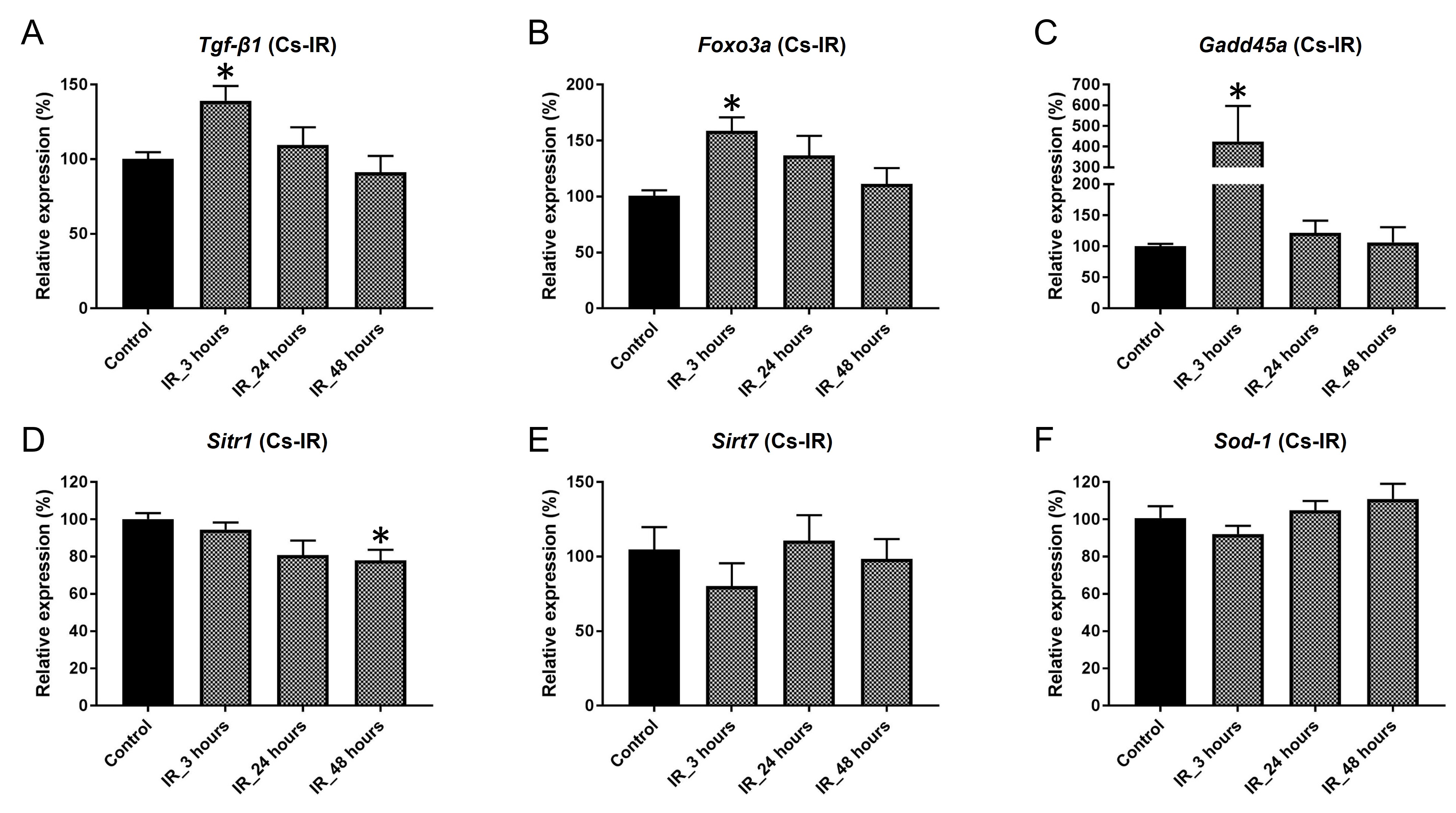
